## Supplementary figures and images for "Contraction-induced endocardial *id2b* plays a dual role in regulating myocardial contractility and valve formation"

### Figure 3-figure supplement 1

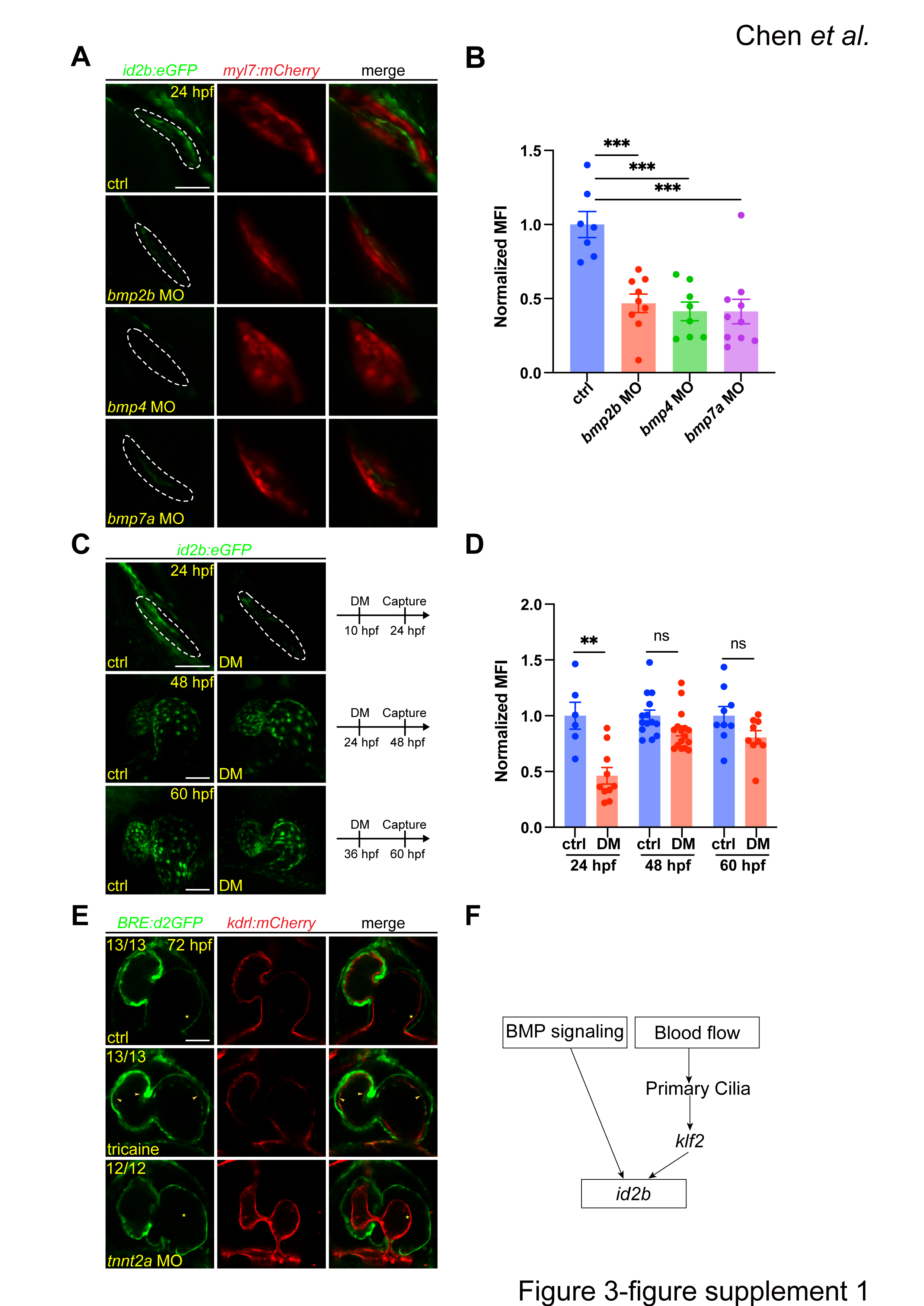

### Figure 4-figure supplement 1

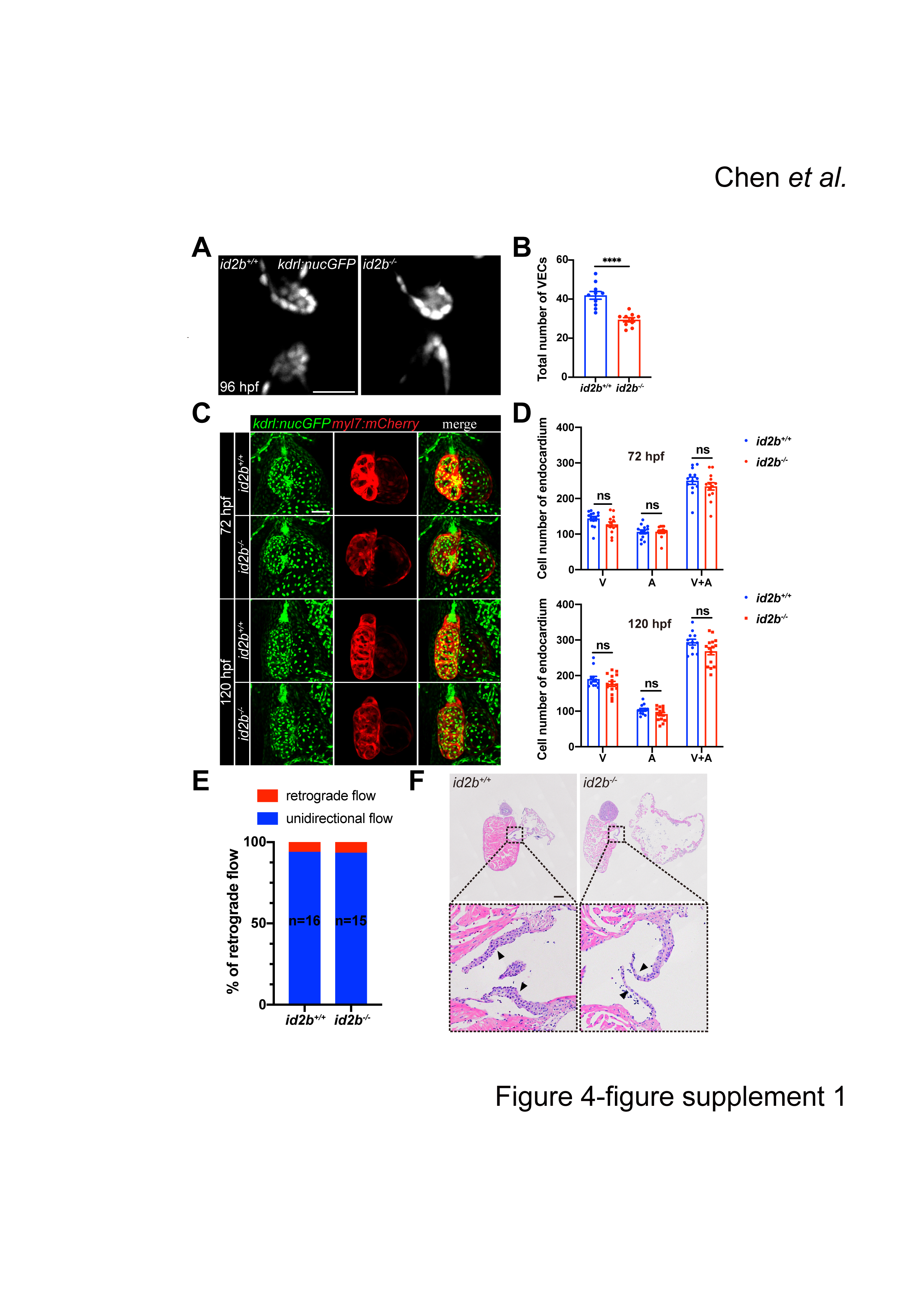

### Figure 5-figure supplement 1

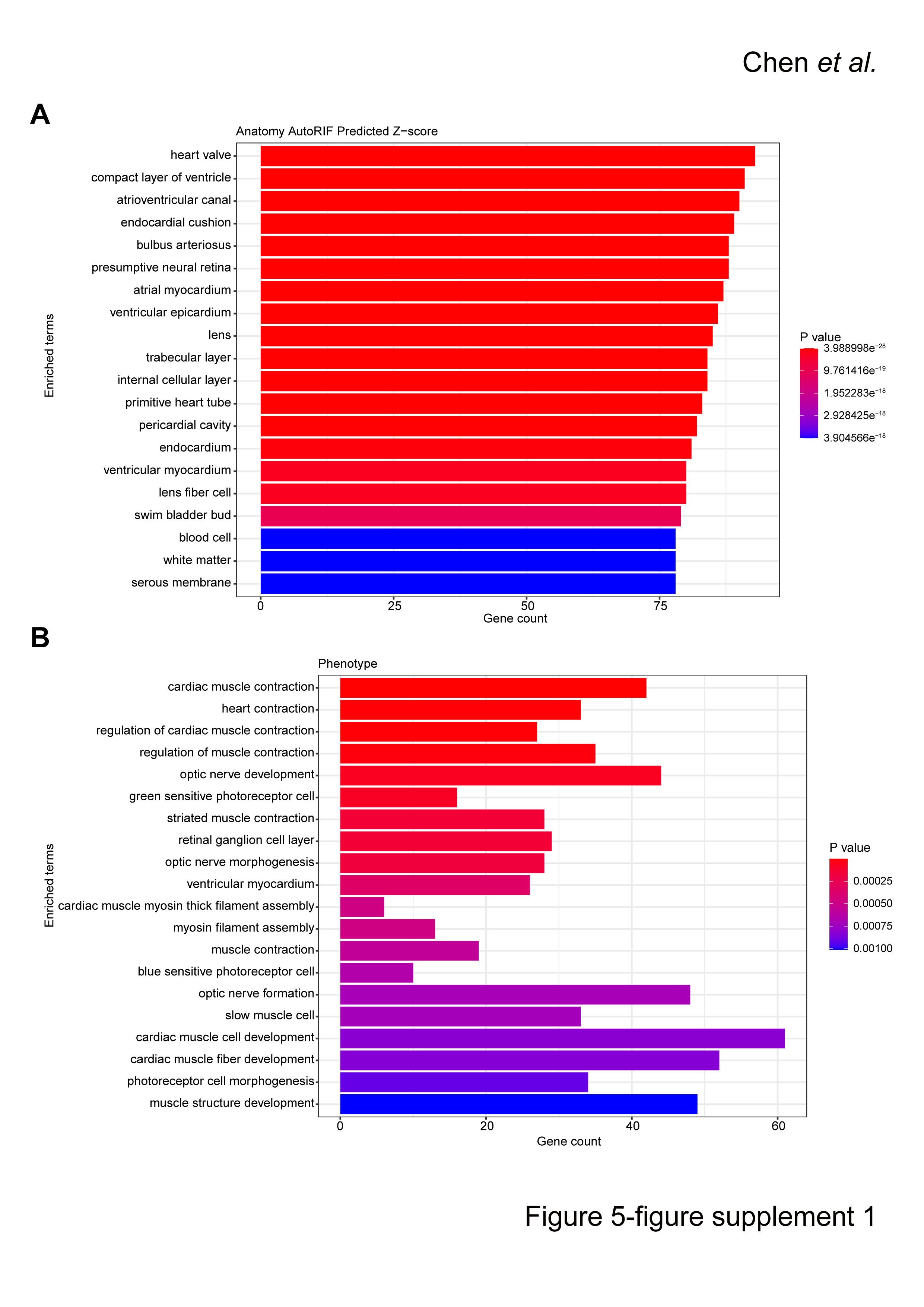

### Figure 5-figure supplement 2

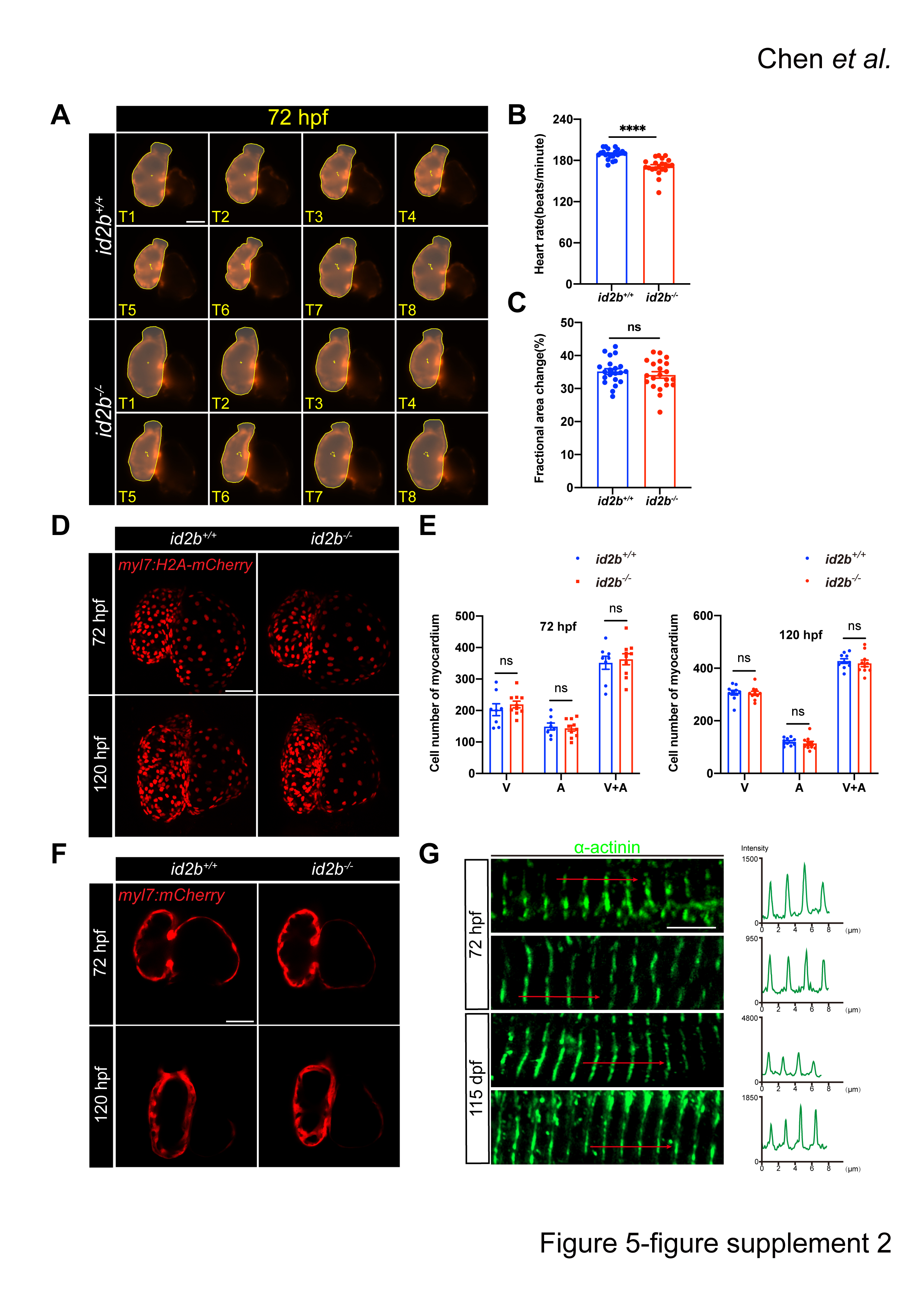

### Figure 6-figure supplement 1

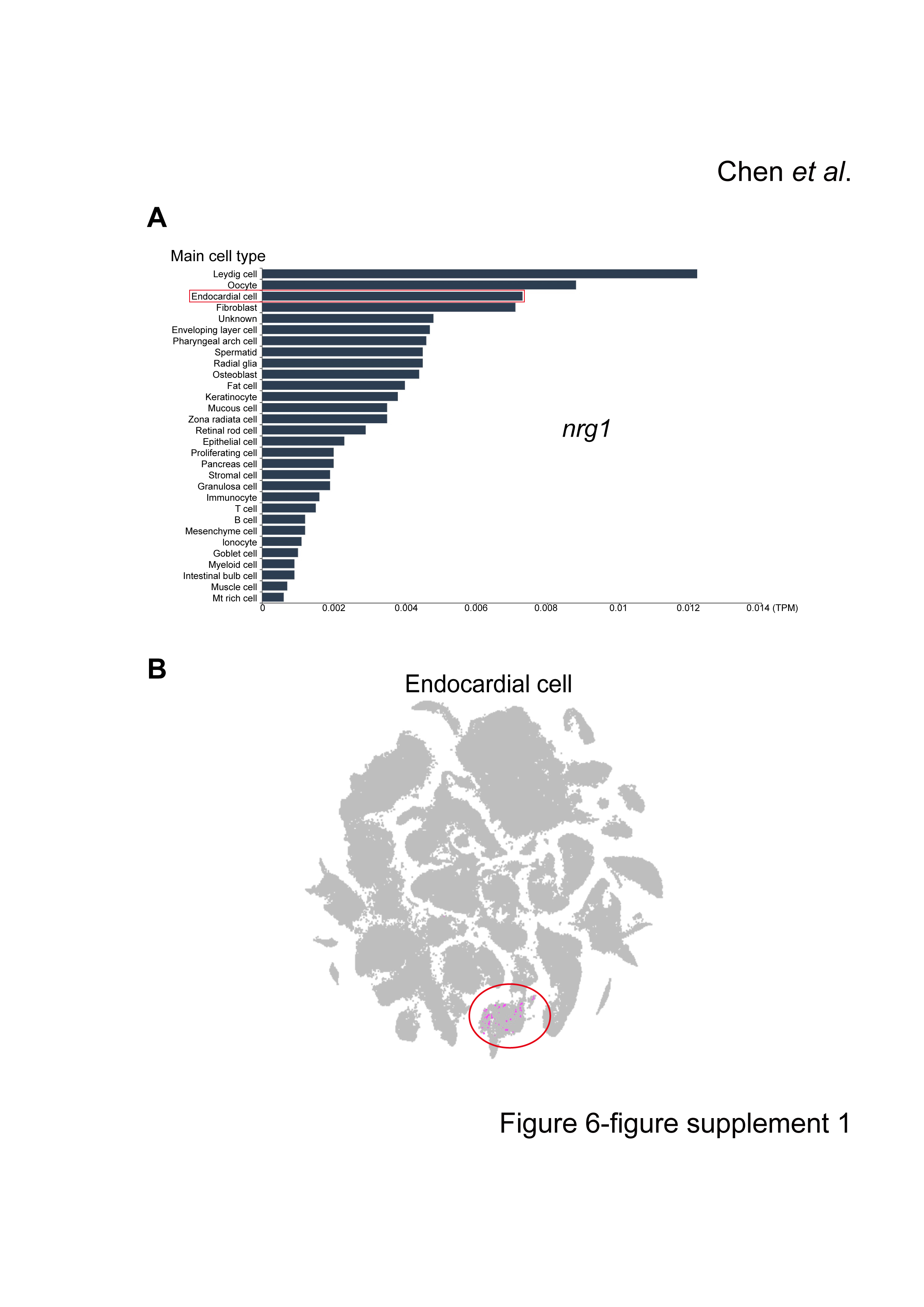

### Figure 7-figure supplement 1

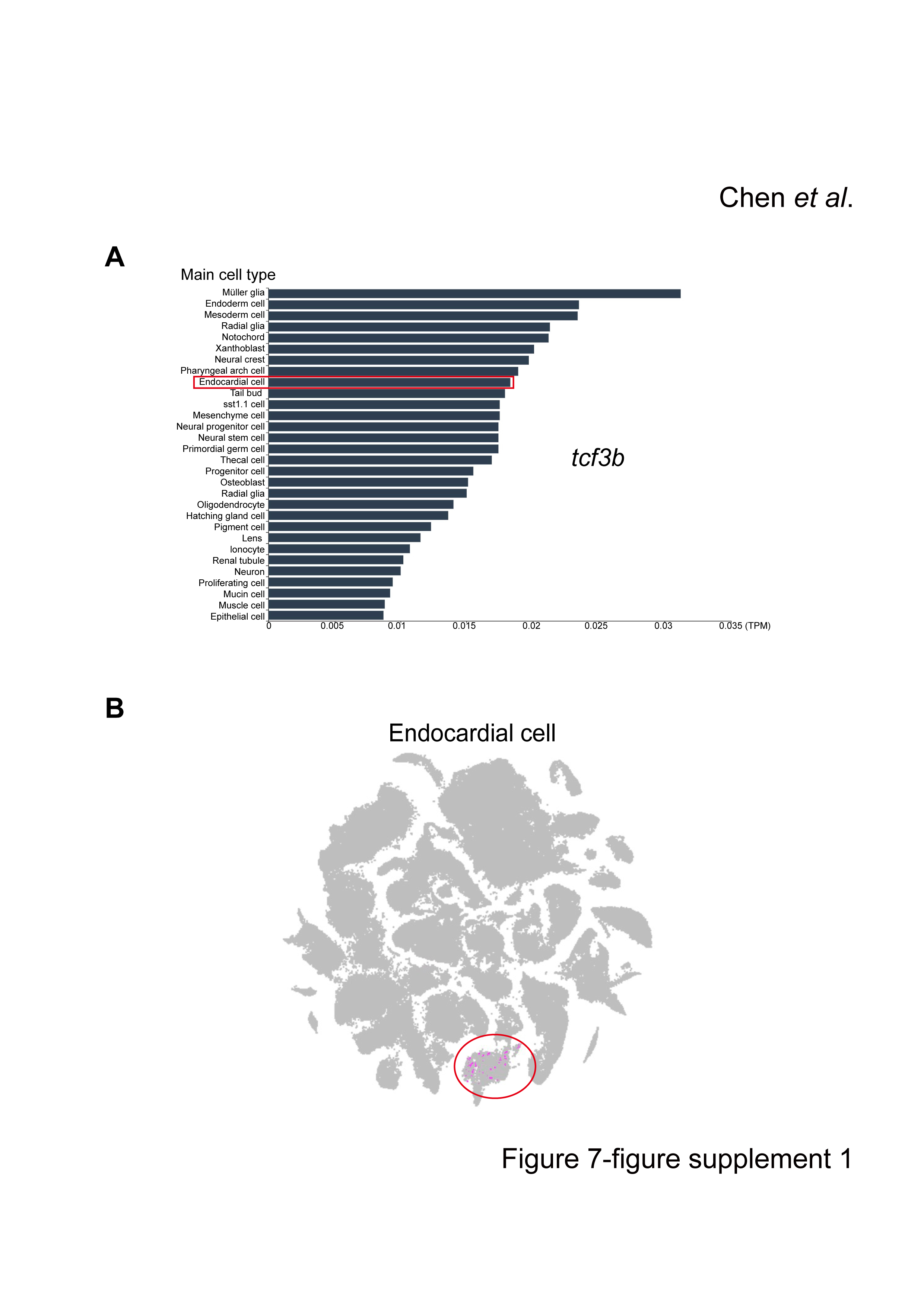
